# Application of Data-Driven computing to patient-specific simulation of brain neuromodulation

**DOI:** 10.1101/2022.09.01.506248

**Authors:** Hossein Salahshoor, Michael Ortiz

## Abstract

We present a class of model-free Data-Driven solvers that effectively enable the utilization of *in situ* and *in vivo* imaging data *directly* in full-scale calculations of the mechanical response of the human brain to sonic and ultrasonic stimulation, entirely bypassing the need for analytical modeling or regression of the data. The well-posedness of the approach and its convergence with respect to data are proven analytically. We demonstrate the approach, including its ability to make detailed spatially-resolved patient-specific predictions of wave patterns, using public-domain MRI images, MRE data and commercially available solid-mechanics software.

## 1 Introduction

Predictive modeling of the mechanical response of the human brain to harmonic stimulation is an important part of many preventive, diagnostic, and therapeutic biomedical procedures [2, 3, 4, 5]. Computational models are increasingly incorporated to inform or guide biomedical research such as transcranial ultrasound stimulation therapies. However, the efficacy of such models depends crucially on precise patient-specific representations of both the geometry and the material response of the brain *in vivo* [6, 7, 8].

In contrast to recent advances in high-resolution, patient-specific, *in-vivo*, and non-invasive imaging techniques, which presently enable accurate representations of brain anatomy down to ultra-fine features [9, 10], the accurate and reliable material modeling of brain tissue has proven challenging and remains a major predictive bottleneck [11, 12]. These modeling challenges arise from the ultra-compliant, complex, heterogeneous, and patient-specific nature of the mechanical properties of brain tissue [12]. These complexities cannot easily be reduced to analytical formulas that differentiate, e.g., between healthy and diseased tissue and the presence of tumors and other lesions in a particular patient.

By way of contrast, recent advances in microscopy and elastography techniques, such as Magnetic Resonance Elastography (MRE) [13], have made possible the *in vivo* characterization of the viscoelastic response of brain tissue in individual patients. Indeed, MRE technology presently allows personalized atlases of brain-tissue viscoelastic properties, including storage and loss moduli as functions of frequency, to be compiled *in vivo* [1, 14]. Whereas present MRE technology covers the sonic frequency range only, it may be expected to eventually encompass the ultrasound range as well. These advances have radically transformed brain biomechanics from a data-starved to a data-rich field, a transformation that challenges theory and computational practice in fundamental and far-reaching ways.

Several possible strategies can be pursued in response to these challenges and opportunities. One currently popular strategy uses supervised Machine Learning (ML) regression in order to fit the data, e.g., using neural-network representations [15]. However, any method of regression, whether based on neural networks or on some other functional representation, is necessarily *biased* and *lossy*. Indeed, regression replaces the empirical data set by a fitted function, a process that inevitably results in loss of information, biases and uncontrolled modeling errors. In addition, models need to be trained *in situ* for specific patients, which is costly and time consuming.

An appealing alternative is to formulate unsupervised, or ‘model-free’, solvers that effectively enable the utilization of *in situ* and *in vivo* imaging data *directly* in calculations, thus entirely bypassing the intermediate material modeling step. In this work, we demonstrate how one such model-free Data-Driven paradigm [16] can be used to forge a direct connection between *in situ* and *in vivo* patient-specific measurements of viscoelastic brain-tissue properties and real-time full-scale calculations of the mechanical response of the human brain to harmonic stimulation.

We ascertain the well-posedness of the proposed Data-Driven solvers analytically and identify physically-motivated properties of the viscoelastic response that ensure convergence with respect to the data. This analysis bears out and complements the numerical convergence tests presented in [16]. For purposes of demonstration, we specifically use a public-domain MRE data set [1] and the corresponding MNI stereotaxic space to develop a finite-element model of the human brain co-registered with the voxel-based MRE data. We then use our model-free Data-Driven solver to compute, directly from the data, wave patterns in the brain resulting from transcranial harmonic stimulation, cf. Section 7.

The unsupervised model-free Data-Driven paradigm demonstrated in this paper suggests a natural path for achieving on-the-fly patient-specific predictions of brain mechanics in a clinical setting. We illustrate the patient-specific aspect of our model-free Data-Driven brain approach by considering MRE data sets from eight different healthy human subjects from the repository provided by [1]. The computed shear strain amplitudes exhibit variations of up to 25%, which, as expected, bears out the need for patient-specific predictions.

We anticipate that strictly unsupervised Data-Driven approaches such as demonstrated here will supply a valuable basis for developing and optimizing patient-specific therapeutic procedures in the future.

## 2 Classical viscoelasticity in the frequency domain

We consider discrete mechanical models comprising *m* material points, or Gauss points in the setting of finite element analysis, whose state at time *t* is characterized by strains *ϵ*(*t*) ≡ {*ϵ*_*e*_(*t*) ∈ ℝ^*d*^, *e* = 1, …, *m*} and stresses *σ*(*t*)≡ {*σ*_*e*_(*t*) ∈ ℝ ^*d*^, *e* = 1, …, *m*}, where *e* labels material points and *d* is the stress and strain dimension. The state of the system is subject to compatibility and dynamic equilibrium constraints of the form

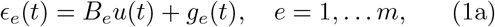

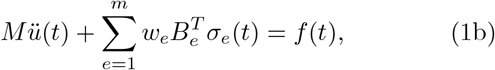

where *u*(*t*) ∈ ℝ^*n*^ is the array of unconstrained nodal displacements, *f* (*t*) ∈ ℝ^*n*^ is the nodal force array, *g*_*e*_(*t*) ∈ ℝ^*d*^ is an initial strain resulting from prescribed displacement boundary conditions, *w*_*e*_ is a positive weight, *B*_*e*_ *∈ L*(ℝ^*n*^, ℝ^*d*^) is the strain-displacement matrix and *M* ∈ ℝ^*n*×*n*^ is the mass matrix. The constraints (1) can also be expressed in compact matrix form as

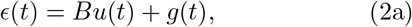

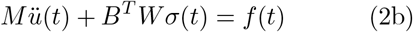

with *B* = (*B*_1_, …, *B*_*m*_)∈ℝ^*N* ×*n*^, *N* = *md, ϵ*(*t*) = (*ϵ*_1_(*t*), …, *ϵ*_*m*_(*t*)) ∈ ℝ^*N*^, *g*(*t*) = (*g*_1_(*t*), …, *g*_*m*_(*t*)) ∈ ℝ^*N*^, *σ*(*t*) = (*σ*_1_(*t*), …, *σ*_*m*_(*t*)) ∈ ℝ^*N*^ and *W* = diag(*w*_1_, …, *w*_*m*_) ∈ ℝ^*N* ×*N*^ is a diagonal matrix with weights *w*_*e*_ as diagonal entries.

Suppose that the forcing is harmonic, i. e.,

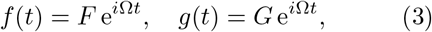

where *F* ∈ ℂ^*n*^ and *G∈*ℂ^*N*^ are constant complex amplitudes and Ω is the frequency of actuation. Alternatively, in Fourier representation

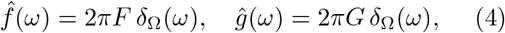

where *δ*_Ω_ denotes the Dirac delta centered at Ω. Harmonic forcing of this type is induced, e. g., as a result of monochromatic transduction. Inserting the representations

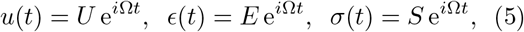

with *U* ∈ ℂ^*n*^, *E∈*ℂ^*N*^ and *S∈*ℂ^*N*^ constant complex amplitudes, the compatibility and dynamic equilibrium equations (2) reduce to the algebraic form

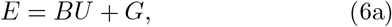

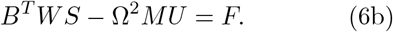

Assume now that all materials in the system are isotropic linear-viscoelastic with complex modulus 𝔼_*e*_(Ω) ∈ℂ, *e* = 1, …, *m*. We recall that the complex moduli have the general form (cf., e. g., [17])

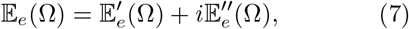

where the real part 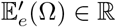 is the storage modulus and the imaginary part 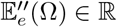 is the loss modulus. The relations

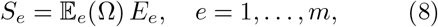

fully characterize the response of the material points in the Fourier representation. We can further collect the relations (8) for all material points into a single matrix relation of the form

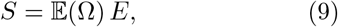

by introducing a diagonal complex-modulus matrix 𝔼 (Ω) = diag(𝔼_1_(Ω), …, 𝔼_*m*_(Ω)) ℂ^*N* ×*N*^ with the complex moduli of all material points as diagonal entries.

Inserting (6a) into (9) and the result into (6b) we obtain the displacement problem

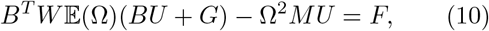

which represents a discrete Helmholtz equation to be solved for the complex displacement amplitude *U* in ℂ^*n*^.

Evidently, the displacement problem (10) has a unique solution if and only if the complex acoustic matrix *B*^*T*^*W* 𝔼 (Ω)*B* − Ω^2^*M* is non-singular, which places restrictions on Ω, *B* and 𝔼 (Ω). It is readily verified that it suffices for the real acoustic matrix *B*^*T*^*W* 𝔼^′^(Ω)*B*−Ω^2^*M* to be non-singular, i.e., that Ω^2^ not be an eigenvalue of *B*^*T*^*W* 𝔼^′^(Ω)*B* with respect to *M* . Thus, in the absence of dissipation solutions only exist away from resonant conditions.

The following proposition shows that dissipation can be relied upon to ensure existence, even under resonant conditions. Here and subsequently, we introduce the norms,

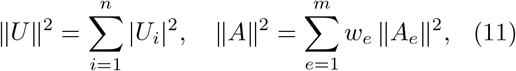

for displacement vectors *U* ∈ ℂ^*n*^ and a local states *A*∈ℂ^*N*^, *N* = *md*, with the choice of norm disambiguated by the argument.

### Proposition 1 (Existence and uniqueness).

*Assume:*

i. *(Material stability) There is a constant c >* 0 *such that, for all e* = 1, …, *m*,

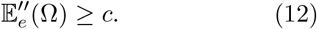
ii. *(Structural stability) There is a constant* 0 *< C <* +∞ *such that, for all U* ∈ ℂ^*n*^,

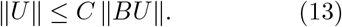

*Then, the complex coustic matrix B*^*T*^*W* 𝔼 (Ω)*B* − Ω^2^*M is non-singular and the displacement problem (10) has a unique solution for all F* ∈ ℂ^*n*^ *and G* ∈ ℂ^*N*^ .

We note that the material stability condition, Prop. 1(i), stipulates a uniform minimum bound for the loss modulus, whereas the structural stability condition, Prop. 1(ii), rules out the existence of mechanisms, i. e., parts of the structure that can undergo motions at zero strain.

## 3 Data-Driven viscoelasticity in the frequency domain

For conventional materials, the complex moduli 𝔼_*e*_(Ω), eq. (8), which completely characterize the viscoelastic behavior of the materials in the structure, can be reliably measured by a number of experimental techniques, including Dynamic Mechanical Analysis (DMA) [18], nanoindentation [19, 20], dynamic shear testing (DST) [21, 22], and others. By contrast, the viscoelastic response of brain tissue exhibits large variations between patients and between different regions of the brain, as well as staggering difficult-to-model complexity [11]. In view of these challenges, we jettison the conventional classical *model-based* paradigm altogether in favor of a *model-free* Data-Driven computational paradigm, which aims to establish a direct connection between patient-specific *in vivo* data and computational prediction.

Following [16], we represent solutions as time histories (*ϵ, σ*) of complex strain and stress, with values (*ϵ*(*t*), ∈ *σ*(*t*)) ∈ ℂ^*N*^ × ℂ^*N*^ at time *t*, in some suitable space 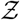 of histories. Specifically, we seek: an admissible history 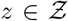 of stress and strain, with value *z*(*t*) ∈ ℂ^*N*^ × ℂ^*N*^ at time *t*, satisfying the field equations (2) at all times; and a material history 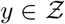 of stress and strain, with value *y*(*t*) ∈ ℂ^*N*^ × ℂ^*N*^ at time *t*, consistent with experimental observation. The objective of the analysis is to determine pairs of histories 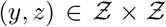 that are closest to each other in the sense of some suitable distance in 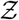, i. e., we seek *y* and *z* such that

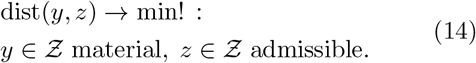

Thus, the Data-Driven solution consists of an admissible history 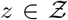 that is closest to being material and material history 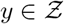 that is closest to being admissible.

For steady problems and harmonic excitation, the stress and strain histories of interest are infinite harmonic ‘wave trains’, and the appropriate space of histories is

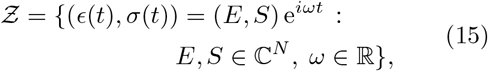

where *E, S* are complex strain and stress arrays, respectively, and *ω* is the frequency. We note that, in the Fourier representation, for every harmonic history 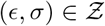, its Fourier transform 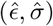 is a Dirac delta in the frequency domain. We also note that 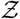 is not a linear space, since it is not closed under addition, but rather a fiber bundle modeled after ℂ^*N*^ × ℂ^*N*^ with based manifold the frequency domain ℝ (cf., e. g., [23]).

In order to explicate problem (14) further, for given forcing *F* ∈ ℂ^*n*^, *G* ∈ ℂ^*N*^ and applied frequency Ω ∈ ℝ we introduce a subspace of 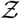 of admissible histories,

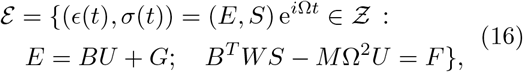

or *admissible set*, consisting of all harmonic stress and strain histories satisfying compatibility and dynamic equilibrium.

Suppose, in addition, that the material behavior is characterized by a material data set 𝒟 that compiles all harmonic histories attainable by the materials, if the material behavior is fully characterized, or a subset of harmonic histories known to be attainable by the material, if the material behavior is partially characterized. Thus, if the complex moduli 𝔼 (*ω*) are fully known for every material point then the material set is

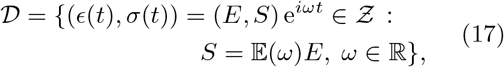

in terms of the complex moduli matrix 𝔼 (*ω*) introduced in (9).

Suppose, contrariwise, that the complex moduli are only partially known through a data set of the form

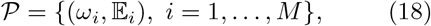

e. g., by testing the materials at *M* frequencies *ω*_*i*_, *i* = 1, …, *M*, and recording the corresponding complex moduli 𝔼_*i*_. In this scenario, the material set is

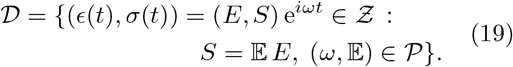

Thus,𝒟 collects all harmonic strain and stress histories consistent with the complex moduli measured at the sampled frequencies, cf. Fig. 2.

**Fig. 1:**
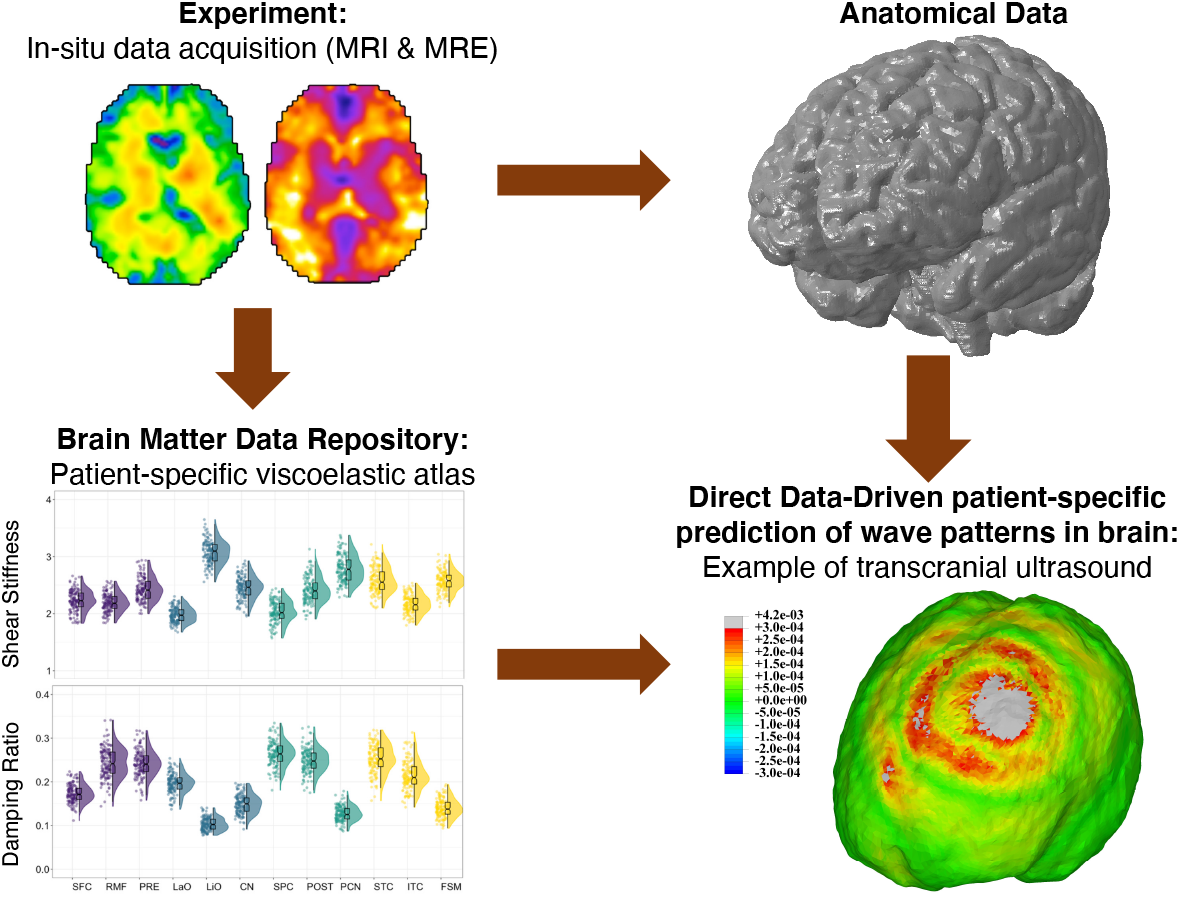
Schematic of the model-free Data-Driven brain mechanics paradigm. Top-left: anatomical data in the form of MRI images [1]. Bottom-left: co-registered atlas of viscoleastic properties of the brain tissue [1]. Top-right: finite-element model of the brain. Bottom-right: steady-state wave pattern induced by lowintensity sonic stimulation of a region on the frontal lobe.

**Fig. 2:**
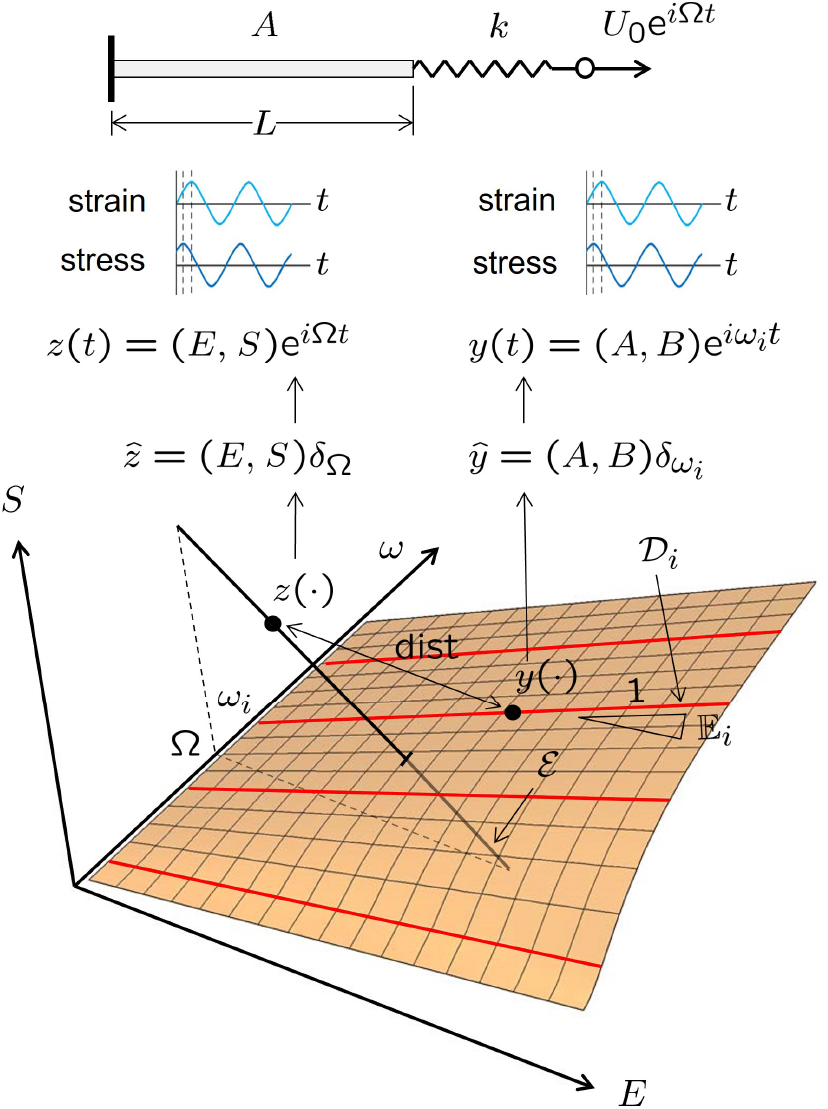
Standard linear viscoelastic bar deforming under the action of harmonic loading imparted through a loading device. Geometrical representation of harmonic histories *z*(*t*) = (*ϵ*(*t*), *σ*(*t*)) = (*E, S*)^*iωt*^ as points in three-dimensional (*E, S, ω*)-space, or *history space* 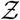. The black line ℰ, the *admissible set*, represents the admissible space of harmonic trajectories satisfying the compatibility and equilibrium equations. The surface 𝒟, *the material set*, encodes the complex moduli 𝔼 (*ω*) for the standard linear solid at different frequencies *ω*. The classical solution is the intersection between the ℰ and 𝒟. In the Data-Driven problem, the complex moduli 𝔼_*i*_ are known only for a sample of *N* test frequencies *ω*_*i*_, *i* = 1, …, *N* . The empirical material set ∈is then the collection of lines 𝒟_*i*_ determined by (*ω*_*i*_, 𝔼_*i*_), shown in red. The Data-Driven solution is the pair of trajectories *y* ∈ 𝒟 and *z* ∈ ℰ closest to each other in some suitable distance.

The Data-Driven problem (14) is now fully defined in the space (15) of harmonic stress and strain histories, with the admissible space (16) encoding all structural geometry and loading data and the material set (17) encoding the known material behavior. Specifically, we wish to determine a pair of harmonic histories 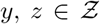 such that

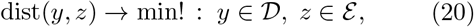

for some appropriate distance in 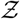. Evidently, the classical solutions are recovered as

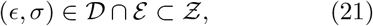

i. e., as the harmonic histories that are both material and admissible, if any. We recall that when the materials are fully characterized and the material set is given by (17), Prop. 1 sets forth condition ensuring the existence of classical solutions.

By contrast, if the material behavior is only partially characterized, the intersection (21) may be empty and classical solutions may fail to exist, whereas the Data-Driven problem (20) does have solutions. The Data-Driven solutions may be regarded as the best possible predictions given the limited information on material behavior.

## 4 A simple illustrative example: Viscoelastic bar

The Data-Driven paradigm just set forth can be visualized and illustrated by means of the simple bar example depicted in Fig. 2. The problem concerns a standard linear viscoelastic bar of cross-sectional area *A* and length *L* deforming under the action of harmonic loading at frequency Ω imparted through a loading device, represented by a spring of stiffness *k*, Fig. 2 (top). The loading device is actuated through a prescribed harmonic displacement *U*_0_e^*i*Ω*t*^. We wish to determine the harmonic stress and strain histories thus induced in the bar.

In this simple scenario, the harmonic histories *z*(*t*) := (*ϵ*(*t*), *σ*(*t*)) = (*E, S*)e^*iωt*^, Fig. 2 (middle), can be visualized as points in three-dimensional (*E, S, ω*)-space, Fig. 2 (bottom), with *E, S* ∈ ℂ and *ω* ∈ ℝ, which supplies a geometrical representation of the space 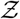 of histories. Considerations of compatibility and equilibrium, including the loading device, give the admissible set

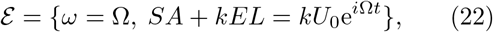

shown as a black line in Fig. 2 (bottom). The complex modulus of the standard linear solid is [16]

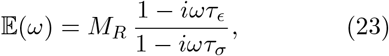

where *M*_*R*_ is the relaxed elastic modulus at zero frequency and *τ*_*ϵ*_ and *τ*_*σ*_ denote strain and stress relaxation times, respectively. We note that the section of the material set 𝒟 at fixed *ω* is a straight line of slope 𝔼(*ω*) in the corresponding (*E, S*)-plane. These lines together define a surface in 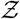 that supplies a geometrical representation of the material set 𝒟, Fig. 2 (bottom). The classical solution is then the unique point of intersection of the admissible line ℰ with the material surface 𝒟 in 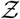.

Suppose, contrariwise, that the complex modulus is only known for a discrete sample of *M* test frequencies *ω*_*i*_, *i* = 1, …, *M*, with 𝔼_*i*_ the value of the complex modulus measured at *ω*_*i*_. Each measured modulus 𝔼_*i*_ determines a line 𝒟_*i*_ in the (*E, S*)-plane at *ω* = *ω*_*i*_, Fig. 2 (bottom). The collection of these *M* lines is the empirical material data set 𝒟, representing the net sum of all empirical knowledge about the material behavior. Suppose that the applied frequency Ω is none of the sampled frequencies *ω*_*i*_. Then, we identify the Data-Driven solution with the pair of histories (*y, z*) with *y* ∈ *𝒟* and *z* ∈ ℰ closest to each other in some appropriate distance defined in 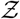, Fig. 2 (bottom).

## 5 Convergence of Data-Driven solutions with respect to the data

Since the admissible and material histories have a Dirac-delta dependence on frequency, a natural notion of distance between histories is supplied by the flat norm [24, 25]. Conveniently, all that is required for present purposes is the flat-norm distance between two Dirac deltas, which has a simple explicit expression. Thus, the flat-norm distance between to Dirac deltas, say *aδ*_*α*_ and *bδ*_*β*_, of amplitudes *a, b* ∈ ℂ^*n*^ and points of application *α, β* ∈ ℝ is, explicitly,

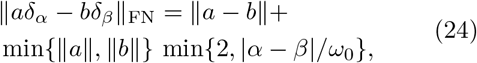

where *ω*_0_ is a reference frequency, Fig. 3a. Thus, the flat-norm distance between two Dirac deltas measures the difference in both amplitude and point of application.

**Fig. 3:**
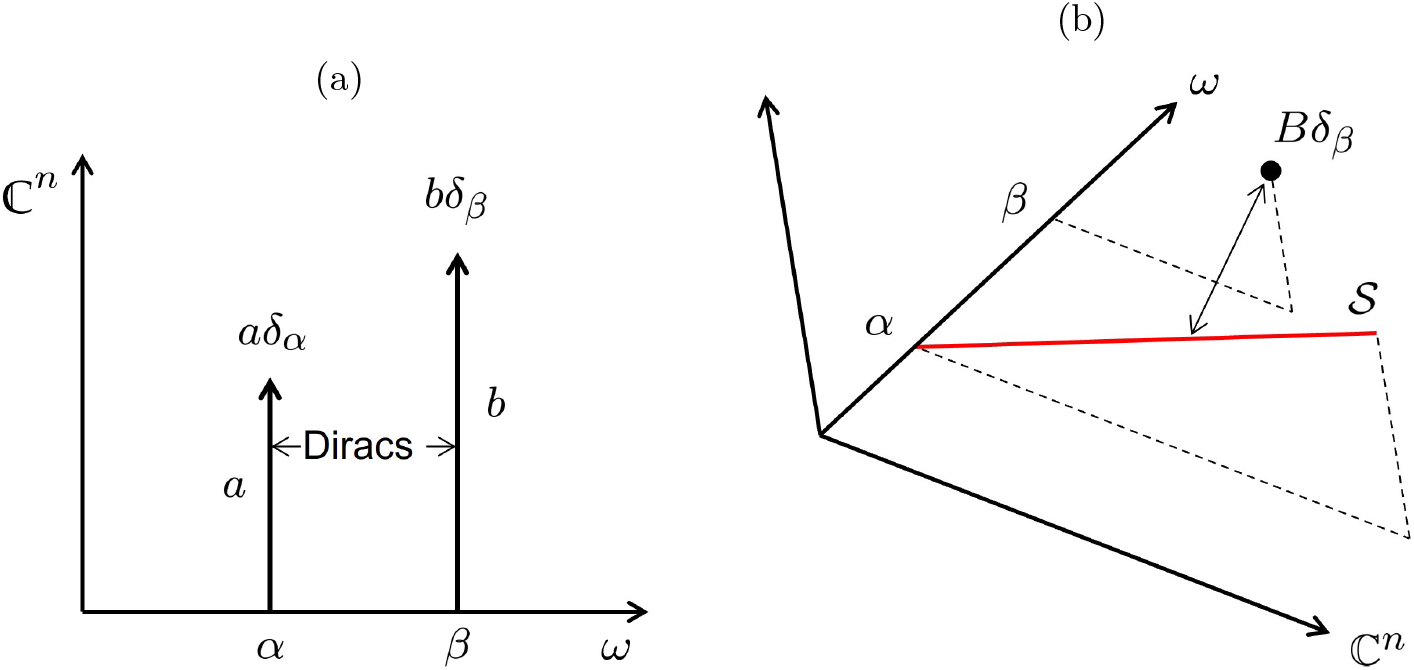
(A) Schematic representation of two Dirac deltas *aδ*_*α*_ and *bδ*_*β*_ of amplitudes *a, b* ∈ ℂ^*n*^ and frequencies *α, β* ∈ ℝ. (B) Computation of distance between a Dirac *Bδ*_*β*_ and a linear subspace 𝒮 of the space of histories 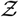.

The flat norm now supplies a natural means of measuring distances between harmonic histories *y* and *z* in 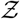, namely,

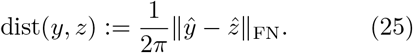

Specifically, if *y* = *Y* e^*iαt*^, *Y* ∈ ℂ^*N*^, and *z* = *Z* e^*iαt*^, *Z* ∈ ℂ^*N*^, we have

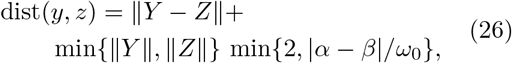

which metrizes the space 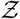 of harmonic stress and strain histories. In applications, we additionally set

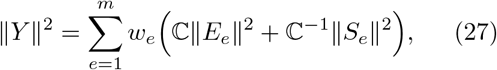

with *Y* = (*E, S*) and ℂ *>* 0 a constant. Thus, the flat-norm distance between two harmonic histories of stress and strain measures the difference in both amplitude and frequency between the histories, as required.

Suppose now that the exact complex moduli 𝔼(*ω*) are not fully known but, instead, for every material point *e* = 1, …, *m*, the material is characterized by a sequence of data sets 𝒫_*h*_, *h* = 1, 2, …, of the form (18) and of increasing size *N*_*h*_ and fidelity along the sequence. Correspondingly, the data sets 𝒫_*h*_ result in a sequence of local material data sets 𝒟_*h*_ of the form (19).

We wish to ascertain conditions under which the corresponding sequence (*y*_*h*_, *z*_*h*_) of Data-Driven solutions converges to the solution (*y, z*), *y* = *z*, of the problem defined by the full— and unknown—complex moduli 𝔼 (*ω*). Convergence with respect to the data of Data-Driven viscoelasticity has been demonstrated in [16] by means of selected verification tests. Here, instead, we seek to characterize the convergence properties of the approximation scheme analytically.

The following proposition sets forth conditions that ensure convergence with respect to the data.

### Proposition 2 (Data convergence).

*Assume that the conditions of Prop. 1 hold and the displacement problem (10) has a unique solution. Let 𝒫*_*h*_ *a sequence of data sets of the form (18). Assume:*

i. *(Lipschitz complex moduli) The complex moduli* 𝔼(*ω*) *are Lipschitz continuous with Lipschitz constant* 𝕃 *>* 0, *i. e*.,

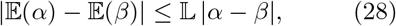

*for all α, β* ∈ ℝ.
ii. *(Bounded complex moduli) The complex moduli are uniformly above and below, i. e*., *there are constants* 0 *<* ℂ_min_ *<* ℂ_max_ *such that*

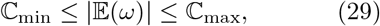

*for all ω* ∈ ℝ.
iii. *(Uniform frequency sampling) There is δ*_*h*_ ↓ 0 *such that, for every h* ∈ ℕ, *there is* (*ω*,𝔼) ∈ 𝒫_*h*_ *with* |*ω* − Ω| ≤ *δ*_*h*_.

*Then, the solution* (*y*_*h*_, *z*_*h*_) *of the* 𝒫_*h*_*–Data-Driven problem converges in the flat norm to the unique solution y* = *z of the* 𝔼(Ω)*–problem*.

We note that assumptions (i) and (ii) of the preceding proposition place regularity and boundedness restrictions on the complex moduli ensuring convergence with respect to the data. The Lipschitz continuity (i) assumption is satisfied, e. g., if 𝔼(*ω*) is differentiable with bounded derivative. The boundedness assumption (ii) requires that the material not loose bearing capacity or become rigid at any frequency. In addition, assumption (iii) requires the data to sample the frequency domain uniformly, in the sense that, for every applied frequency Ω, there exists a sufficiently close empirically sampled frequency with sampling error decreasing to zero along the data set sequence.

## 6 The Data-Driven viscoelastic solver

Algorithm 1 summarizes a Data-Driven solver that seeks to minimize distance between admissible and material histories by means of a fixed point iteration. Conveniently, the solver can be implemented as a wrapper around conventional FE element programs, including commercially available software. Further details of implementation, such as the computation of the flat norm, and tests of convergence of the fixed-point iteration and of the solution with respect to the data, may be found in [16].

For purposes of choosing the frequency *ω* among those sampled experimentally that best matches a trial admissible state *z* at the applied frequency Ω, it becomes necessary to compute the distance between *z* and the empirical material subspaces

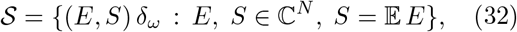

with (*ω*,𝔼) ∈ 𝒫, cf. Figs. 2 and 3b. The general problem of determining the flat-norm distance between a Dirac *Bδ*_*β*_ and a linear subspace *𝒮* of 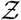 has been addressed in [16]. The result is, explicitly,

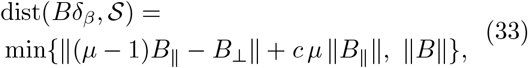

with

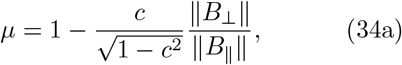

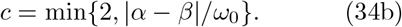

In (33), we assume the norms in (27) to be Euclidean and *B*_∥_ and *B*_⊥_ denote the parallel and perpendicular components of *B* with respect to *𝒮*.

Apart from being natural for infinite wave trains such as harmonic histories, the flat norm distance affords the great practical advantage of enabling the use of data at frequencies other that the applied frequencies of excitation, with an appropriate penalty accorded to the discrepancy between the two. This property is specially advantageous when the data is sampled at discrete frequencies, which are likely to differ from the applied frequency.

It bears emphasis that the Data-Driven paradigm adhered to in this work is strictly unsupervised in that the entire material data set—and nothing but the material data set—is used directly in the calculations without modeling, regression or reduction of any type.

### Algorithm 1 Data-driven solver, harmonic loading.

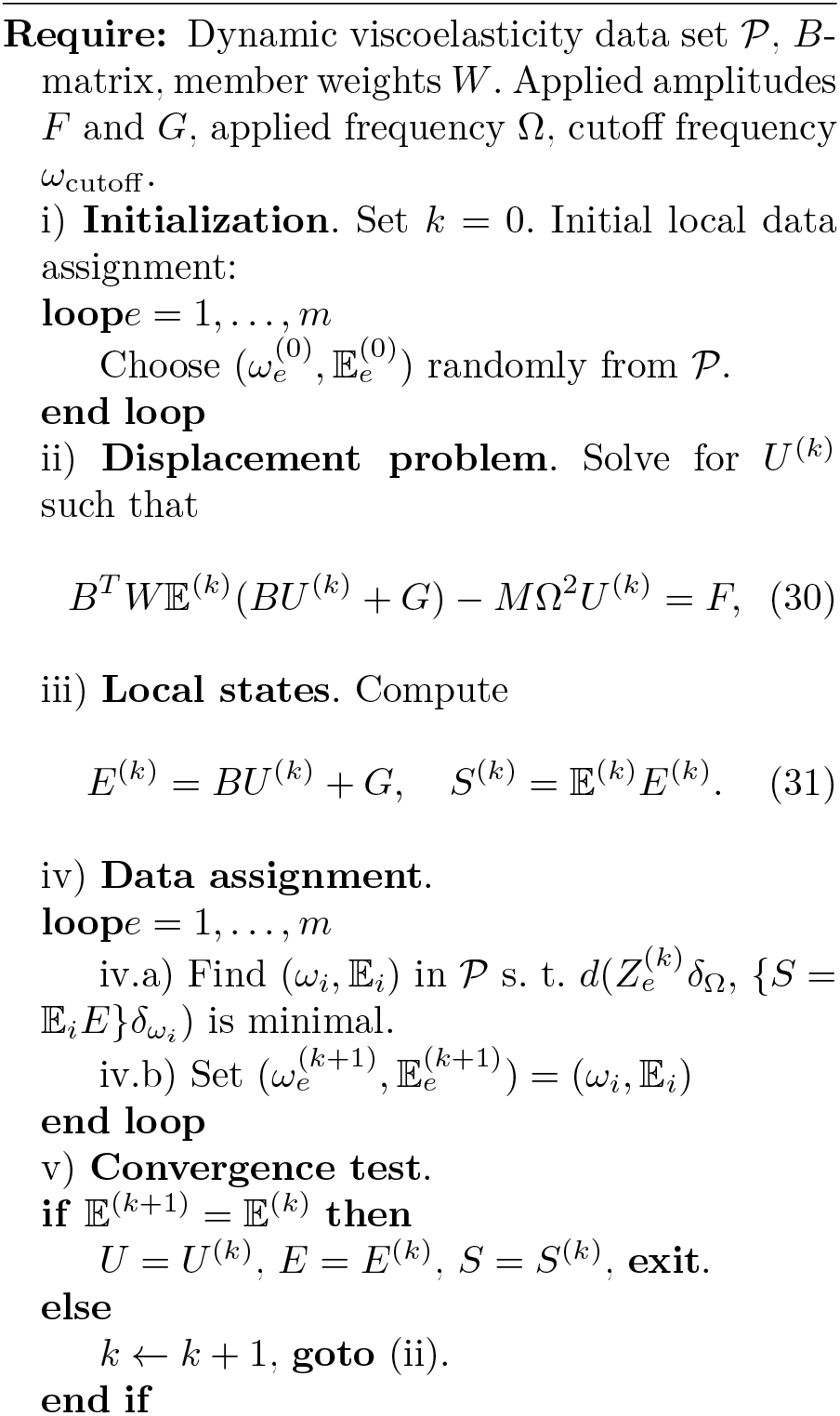

## 7 Application to transcranial acoustic neuromodulation

A flowchart of the model-free Data-Driven paradigm presented in the foregoing, as it bears on transcranial sonic stimulation, is shown in Fig. 1. We use anatomical data, in the form of MRI images, Fig. 1 top-left, in conjunction with a co-registered atlas of viscoleastic properties of the brain tissue, Fig. 1 bottom-left, to generate a finite-element model of the brain, Fig. 1 top-right. We choose the finite-element mesh size equal to, or smaller than, the voxel size in the MRE atlas, so that every material point in the mesh can be assigned to a unique MRE voxel. We then solve in the frequency domain for the steady-state wave pattern induced by sonic transduction, Fig. 1 bottom-right.

### 7.1 Data-Drivencalculationsbasedonaverageproperties

In the present work, we use the recently published atlas of viscoelastic properties of brain tissue [1], acquired at a single frequency of 50 Hz and averaged over 134 human subjects. We also use the associated MRI coordinate data from the same source to develop a computational heterogeneous brain model, co-registered to the MRE data. The resultant geometry is discretized into a finite element model comprising 0.2 million three-dimensional tetrahedral elements, Fig. 4a.

**Fig. 4:**
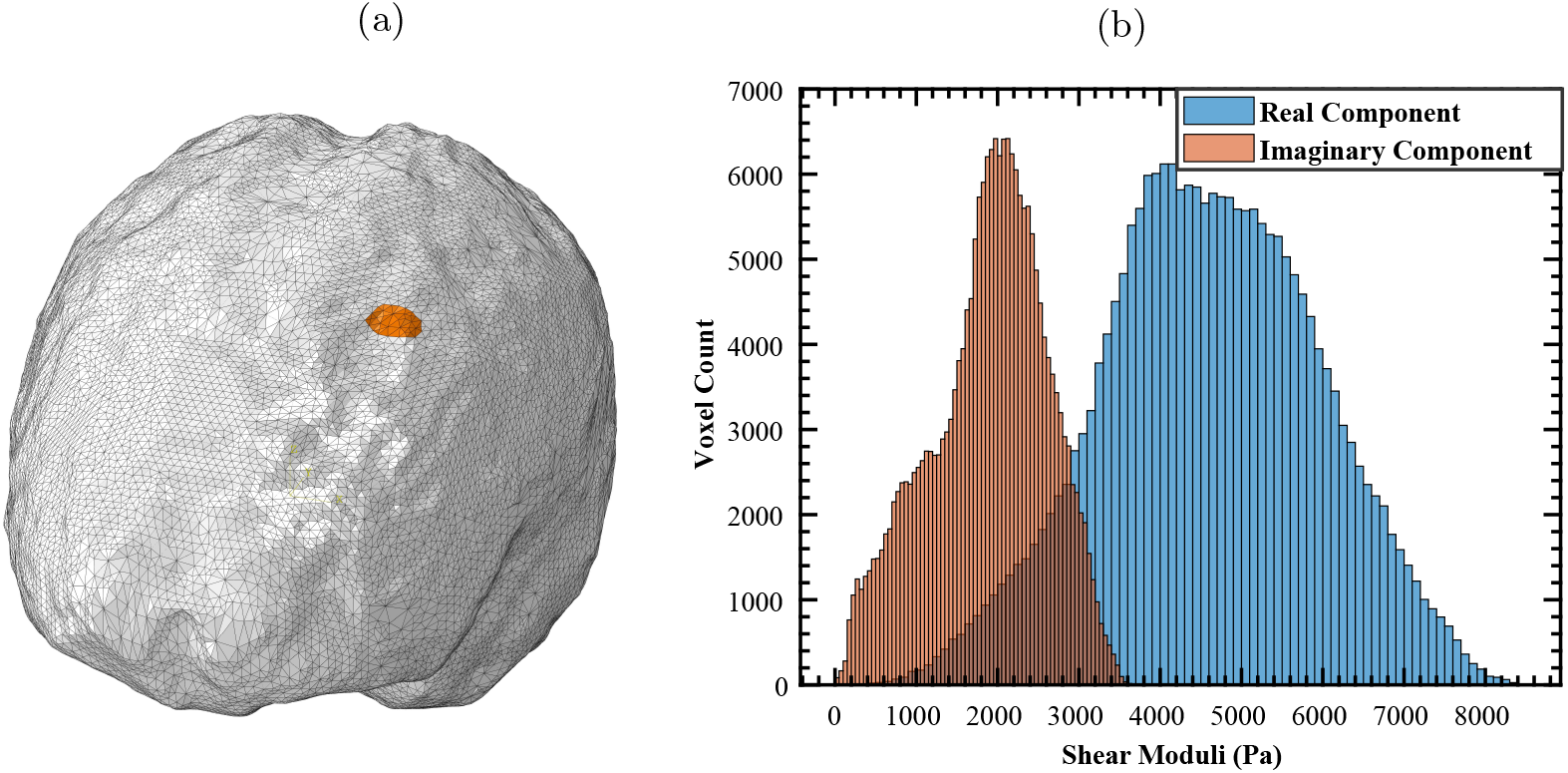
(A): Finite element model of a heterogeneous brain with 0.2 million tetrahedral elements. Transcranial stimulation is modeled by subjecting the highlighted region to harmonic pressure as a traction boundary condition. (B): histogram of complex shear moduli from MRE data. The computational model is co-registered to the data. Both the geometry and viscoelastic map are obtained from public domain MRE data [1].

We note that the MRE data encodes complex viscoelastic shear moduli with real and imaginary components, corresponding to storage and loss moduli for each material point, respectively. For the sake of simplicity, we assume the brain matter to be isotropic, with an elastic bulk modulus of 2 GPa assigned to the entire model. We remark that, conditioned on the availability of the data, the present framework can account for fully anisotropic brain matter or other material symmetries thereof.

Fig. 4b shows a histogram of the complex moduli in the material data set binned by voxel. The wide spread of properties in the material data set resulting from material heterogeneity, patient-to-patient variability and measurement scatter is particularly striking.

In calculations, we subject a small region on the frontal lobe of the brain, shown in orange in Fig. 4, to a harmonic pressure with a peak amplitude of 60 kPa. Calculations were performed using the commercial finite-element software Abaqus (Abaqus/Standard, Dassault Systemes Simulia, France) as compute engine within the model-free Data-Driven wrapper.

Fig. 8 shows computed contours of the real and imaginary components of all six components of the strain tensor across the entire brain. The relatively higher magnitude of the shear strains with respect to volumetric strains is expected from the softness of the brain tissue in shear.

The relative ease of implementation of the framework on top of readily available commercial software and the speed of the calculations, comparable to that of a simple elastic analysis, is noteworthy, as is the fully unsupervised Data-Driven nature of the calculations, which follow directly from *in vivo* data and entirely bypass the traditional material modeling step of computational mechanics.

### 7.2 Patient-specificData-Drivencalculations

We illustrate the patient-specific scope of the model-free Data-Driven paradigm by recourse to MRE data sets from eight different healthy human subjects from the repository provided by [1]. The subjects were four male and four female with ages varying from 18 to 35. As in the preceding section, we solve the corresponding eight Data-Driven problems with all components of the computational model held fixed except for the complex moduli data. Histograms of the material data set binned by voxel for all eight patients are depicted in the supplementary material. We again emphasize that all calculations are unsupervised and based directly on the MRE data without recourse to analytical modeling of the data at any stage of the calculations.

Fig. 6 shows the computed maximum complex amplitude of harmonic shear strains at a frequency of 50 Hz for all eight patients. Computed contours of real and imaginary components of both maximum and minimum principal strain across the entire brain for all eight patients are shown in the supplementary material. We observe that the real component of maximum shear strain *ϵ*_*xy*_ varies from 0.015 to 0.021, or a 25% spread, with similar variations for the remaining components. These results bear out the importance of patient-specific data and illustrate the potential of the model-free Data-Driven paradigm for achieving patient specificity on demand in a clinical environment.

**Fig. 5:**
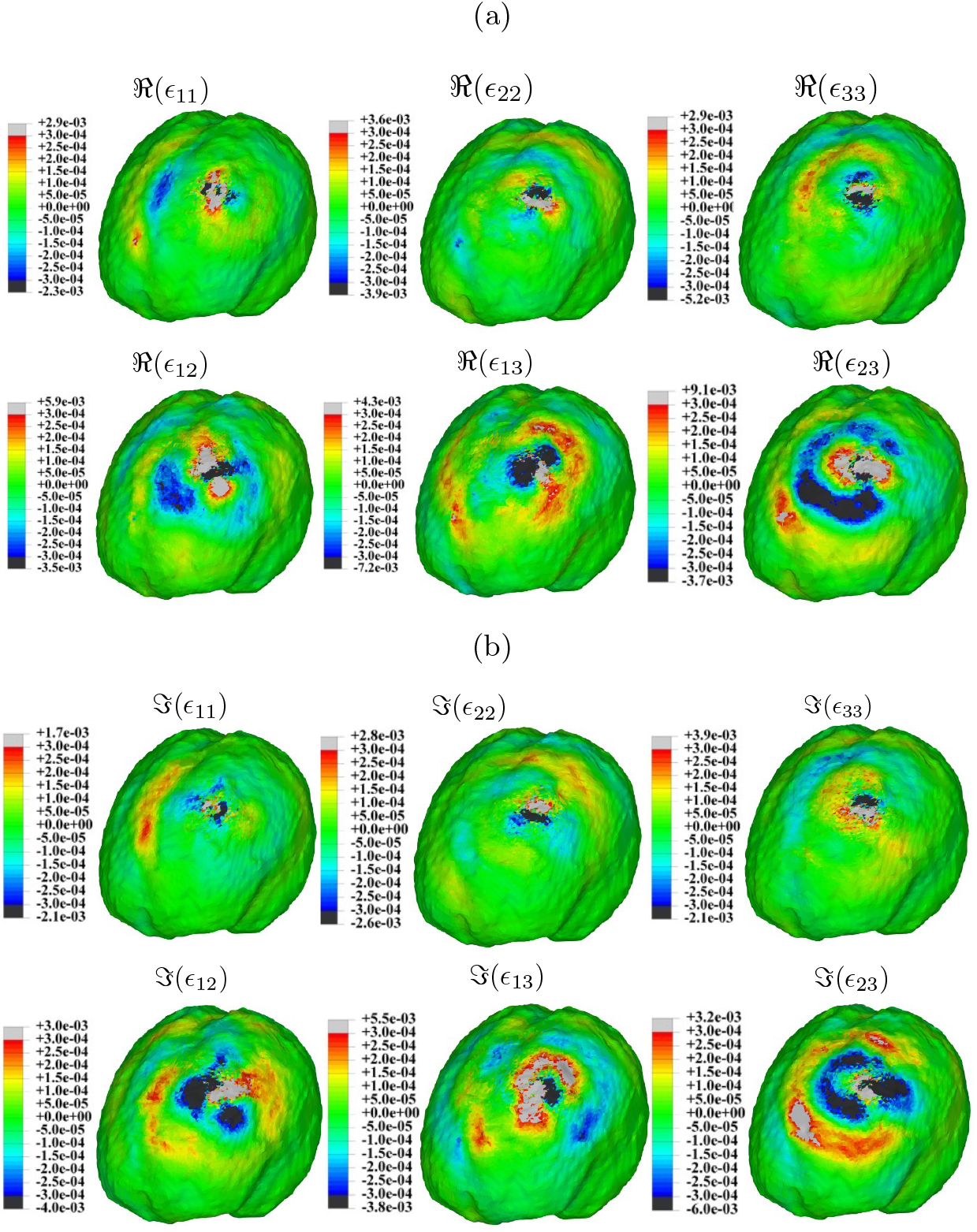
Model-free Data-Driven simulation of transcranial sonic stimulation on a region on the frontal lobe of brain, Fig. 4a. Spatial distribution of components of strain at steady state. a) Real part; b) Imaginary part.

**Fig. 6:**
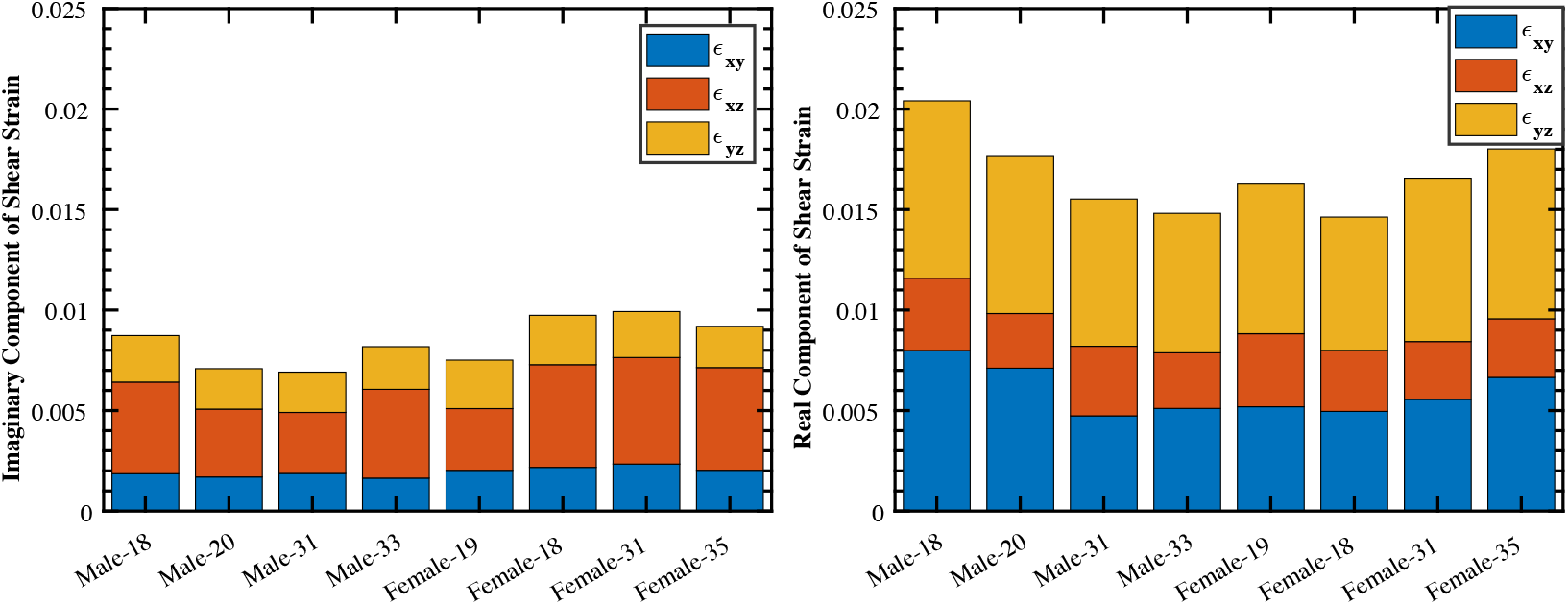
Model-free Data-Driven simulation of transcranial sonic stimulation on a region on the frontal lobe of brain, Fig. 4a, for eight patient-specific MRE data sets. Maximum strain amplitudes at steady state. Left: Imaginary part; Right: Real part.

### 7.3 Data-Drivencalculationsintheultrasoundrange

Data from *in vivo* MRE are presently limited to low frequency and does not cover the ultrasonic range [1, 14]. However, given current developments in elastography [26], as well as in other emerging full-field microscopy technologies [27], experimentally measured viscoelastic atlases at ultrasound frequencies may be expected to become available in the foreseeable future.

In anticipation of these developments, we proceed to demonstrate that the proposed Data-Driven solver is applicable to ultrasound neuromodulation (UNM). To that end, we synthesize a set of spatially-varying atlases of complex shear moduli for frequencies in the range of 90 kHz to 110 kHz. The data samples a standard linear model with complex shear modulus calibrated from brain matter by [28], Fig. 7.

**Fig. 7:**
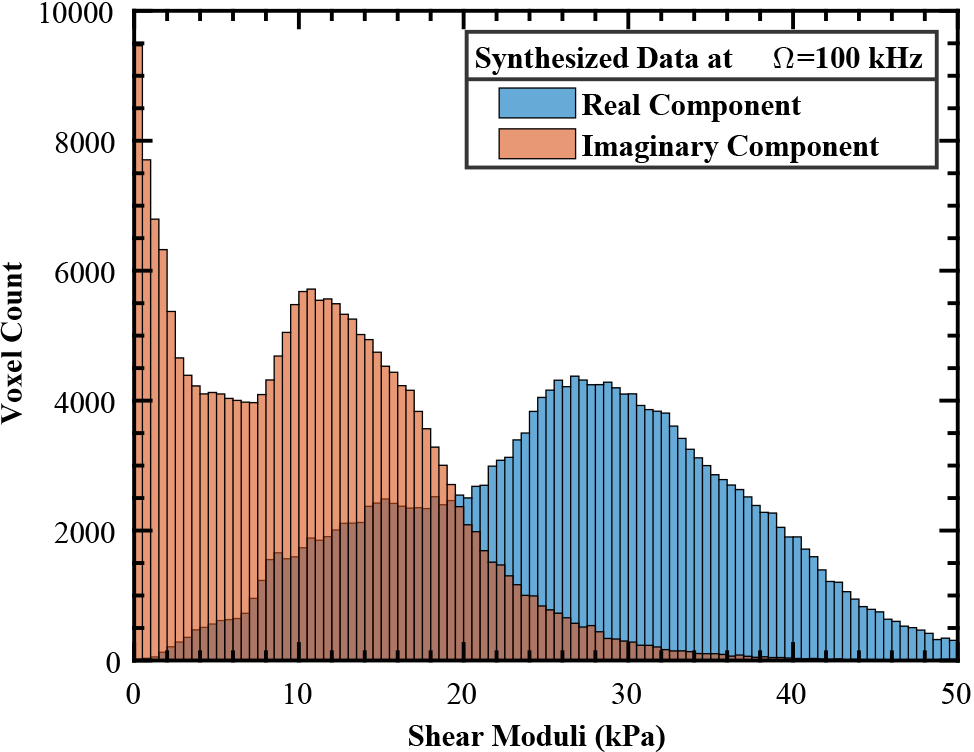
Histogram of synthesized complex shear moduli data at 100 kHz, representative of the data utilized for ultrasonic neuromodulation simulations.

We again subject a region on the frontal lobe of the brain, shown in orange in Fig. 4, to a harmonic pressure with a peak amplitude of 100 kPa and central frequency of 100 kHz in the ultrasound range, corresponding to low-intensity focused ultrasound (LIFUS) stimulation. The high-frequency range is computationally challenging because it sets stringent spatial resolution requirements concomitant to the short wavelength of the induced wave patterns. A convergence study showing that the resolution of the mesh used in the calculations meets these resolution requirements has been presented in [5].

Calculations were again performed using the commercial finite-element software Abaqus (Abaqus/Standard, Dassault Systemes Simulia, France) as compute engine within the model-free Data-Driven wrapper. Fig. 8 shows computed contours of the real and imaginary components of all components of the strain tensor across the entire brain. As may be seen from the figure, despite the peak pressure increase from 60 kPa to 100 kPa, the maximum shear strains exhibit a modest decrease from 9.1 *×* 10^−3^ to 5.7 *×* 10^−3^ relative to the sonic case, which showcases the sensitivity of the results to the viscoelastic response of the brain tissue as a function of frequency. The strain fields in Fig. 8 are also strikingly different from those in Fig. 6, corresponding to the sonic range, in that the ultrasonic fields are much more localized that the sonic fields and exhibit patterning on a finer scale. These differences are expected from the dispersion relation of the material and evince the need for fine spatial resolution in the ultrasound range.

**Fig. 8:**
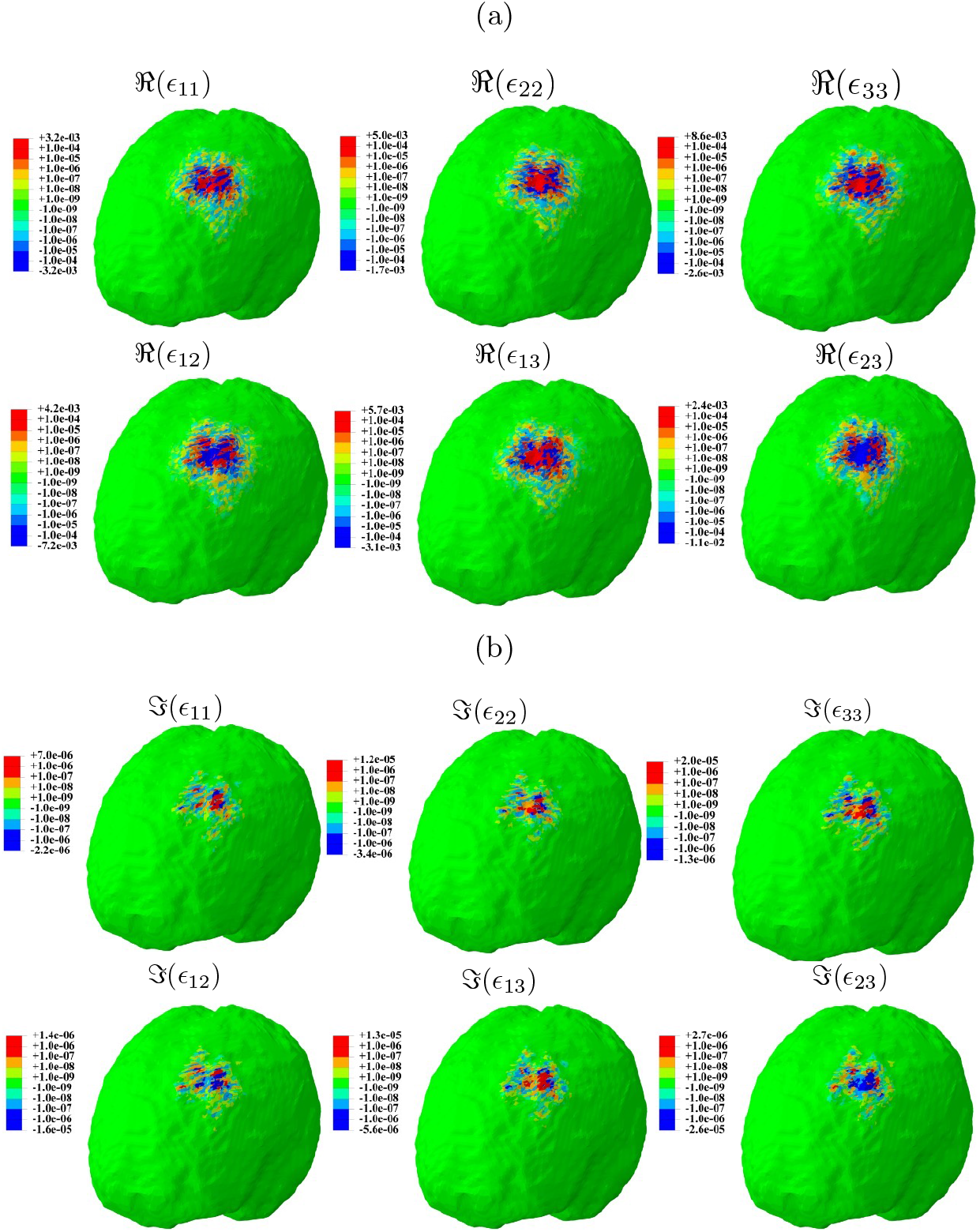
Model-free Data-Driven simulation of ultrasonic neuromodulation with peak pressure of 100 kPa and central frequency of 100 kHz, Fig. 4a. Spatial distribution of components of strain at steady state. a) Real part; b) Imaginary part.

## 8 Summary and concluding remarks

We have presented a class of model-free Data-Driven solvers that open the way for the utilization of *in situ* and *in vivo* imaging data *directly* in full-scale calculations of the mechanical response of the human brain to sonic and ultrasonic stimulation, entirely bypassing the need for analytical modeling or regression of the data. Thus, the specific Data-Driven solver presented in this work falls within the general strategy of unsupervised set-oriented machine learning (cf., e. g., [29]).

The ability to make predictions directly and on-the-fly from clinical data is important due to the variability of the data from patient to patient, between different regions of the brain for a given patient, and the complexity of the frequency-dependent viscoelastic response of brain tissue, which challenge traditional modeling and calibration (cf. [11] for a striking account).

The well-posedness of the approach and its convergence with respect to data are proven analytically. In particular, we uncover physically-motivated restrictions on the complex moduli that ensure convergence to the ground-truth behavior as the resolution provided of the data set is increased. This convergence analysis thus bears out and complements the numerical tests presented in [16].

We have demonstrated the range and scope proposed Data-Driven solver, including its ability to make detailed spatially-resolved patient-specific predictions of wave patterns, using public-domain MRI images, MRE data and commercially available solid-mechanics software (Abaqus/Standard, Dassault Systemes Simulia, France) in simulations of low-intensity sonic and ultrasonic brain stimulation. The simulations bear out the importance of patient-specific data and illustrate the potential of the model-free Data-Driven paradigm for achieving patient specificity on demand in a clinical setting.

The opportunity for a paradigm shift to data-centric simulation owes fundamentally to remarkable recent advances in microscopy and elastography techniques, such as Magnetic Resonance Elastography (MRE) [13], that have made possible the *in vivo* characterization of the viscoelastic response of brain tissue in individual patients. These advances have effectively transformed computational brain mechanics from a data-poor to a data-rich field. This transition in turn raises heretofore unprecedented opportunities but also challenges the theory and praxis of scientific computing in ways that demand extensive further research.

## Acknowledgments

This project was supported by U.S. National Institutes of Health Grant No. 1RF1MH117080 and by the German Research Foundation (Deutsche Forschungsgemeinschaft; DFG) within the Priority Program 2311, grant number 465194077.

## A Proofs of theorems

### Proof of Prop. 1.

Suppose there is *U* ∈ ℂ^*n*^ such that

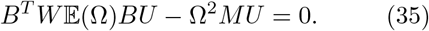

Testing this equation with *U*, we obtain

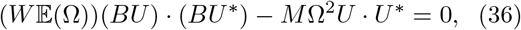

and taking the imaginary part,

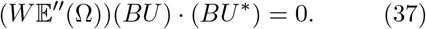

By assumptions (i) and (ii),

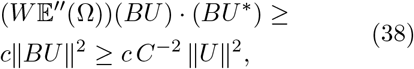

which, together with (37) implies *U* = 0, as required. □

### Lemma 3

*Let ω*_0_ *>* 0; *α, β* ∈ ℝ, |*β* − *α*| ≤ 2*ω*_0_; 𝔼(*α*), 𝔼(*β*) ∈ ℂ. *Let E, S* ∈ ℂ^*N*^, *S =* 𝔼 (*β*) *E, Z* = (*E, S*), 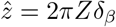. *Let*

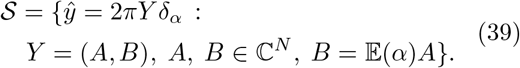

*Assume:*

i. *The complex modulus* 𝔼(*ω*) *is Lipschitz continuous with Lipschitz constant* 𝕃 > 0, *i. e*.,

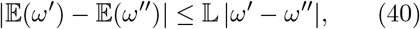

*for all ω*^*′*^, *ω*^*′′*^ ∈ ℝ.
ii. *The complex modulus is bounded above and below, i. e*., *there are* 0 *<* ℂ_min_ *<* ℂ_max_ *such that*

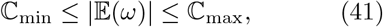

*for all ω* ∈ ℝ.

*Then, there is C >* 0 *such that*

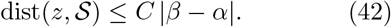

*Proof*. Let *Y* = (*A, B*) be the closest-point projection of *Z* on the linear subspace {*B* = 𝔼(*α*)*A*}. A straightforward calculation gives

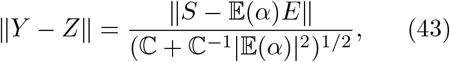

and

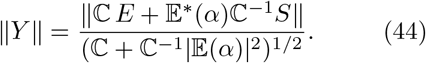

By orthogonality, ∥ *Y* ∥ ≤ ∥ *Z* ∥ and, from (24),

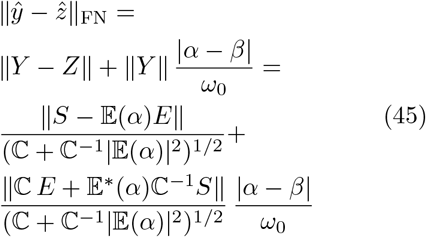

Assume *S* = 𝔼 (*β*)*E*. Then,

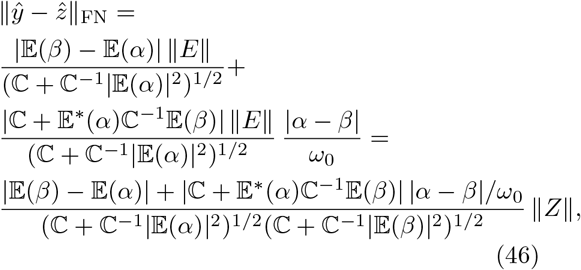

By (i),

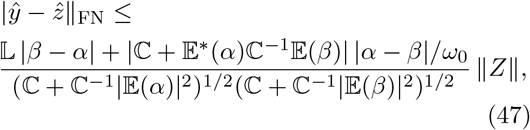

and by (ii),

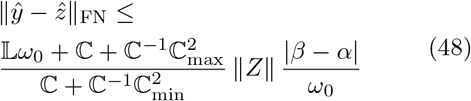

and (42) follows with

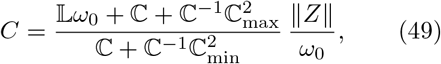

as advertised. □

### Lemma 4 (Transversality).

*Suppose that the matrix B*^*T*^*W*𝔼(Ω)*B is non-singular and the viscoelasticity problem (10) has a unique solution for every* Ω ∈ ℝ, *F* ∈ ℝ^*n*^ *and G* ∈ ℝ^*N*^ . *Let* 𝒟 *be as in (17) and ε as in (16). Assume assumption (ii) of Lemma 3 holds and choose the constant* ℂ *in (27) such that* ℂ_min_ ≤ ℂ ≤ ℂ_max_. *Then, there are c >* 0, *b* ≥ 0 *such that*

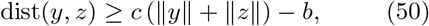

*for all y* ∈ 𝒟 *and z* ∈ ℰ.

*Proof*. The intersection of *𝒟* and *ε* consists of one single point *z*_0_ corresponding to the unique solution of the linear viscoelasticity problem (10). Let ℰ_0_ be the constraint set for zero forcing, i. e., (16) with *F* = 0 and *G* = 0. Then, by linearity, we have ℰ = *z*_0_ + ℰ_0_. We claim that there is *c* > 0 such that

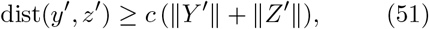

for every *y*^*′*^ = *Y* ^*′*^e^*iαt*^ ∈ 𝒟 and *z*^*′*^ = *Z*^*′*^e^*iβt*^ ∈ *ℰ*_0_. Write *Z*^*′*^ = (*E*^*′*^, *F*^*′*^) ∈ ℂ^*N*^ *×* ℂ^*N*^ . Then,

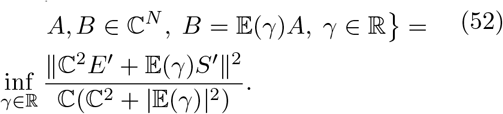

Expanding the square with *z*^*′*^ ∈ ℰ_0_, we obtain

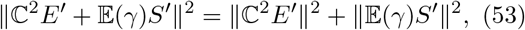

whence

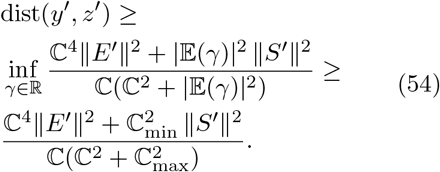

With ℂ_min_ ≤ ℂ ≤ ℂ_max_, we further have

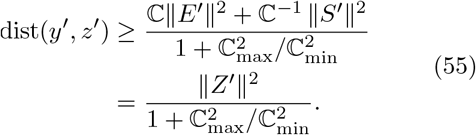

Finally, the estimates

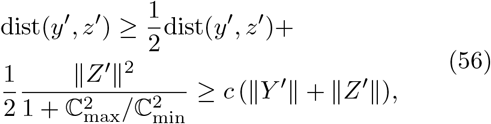

yield (51), as surmised. Let now *y* ∈ 𝒟, *z* ∈ ℰ, and define *z*^*′*^ = *z* − *z*_0_ ∈ ℰ_0_. Then a triangular inequality gives ∥*Z*^*′*^∥ ≥ ∥*Z*∥−∥*Z*_0_∥ and we obtain

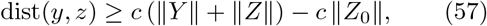

which gives (50) with *b* = *c* ∥*Z*_0_∥. □

### Proof of Prop. 2.

Let *y*_∗_ = *z*_∗_ be the unique solution of the 𝔼 (Ω)–problem. Set 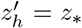 and let 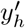 the closest point to *z*_∗_ on the subspace

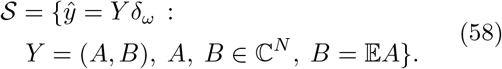

with (*ω*,𝔼) ∈ 𝒫_*h*_ such that |*ω*− Ω| ≤ *δ*_*h*_. Then, by optimality,

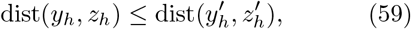

where (*y*_*h*_, *z*_*h*_) is the solution of the 𝒫_*h*_–Data-Driven problem. By Lemma 3,

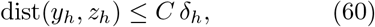

whence in follows that dist(*y*_*h*_, *z*_*h*_) → 0. By Lemma 4, it follows that, possibly up to subsequences, there exist *Y, Z* ∈ C^*N*^ such that *Y*_*h*_ → *Y* and *Z*_*h*_ → *Z*. By the continuity of the distance, it follows that dist(*y, z*) = 0, hence *y* = *z* is classical solution. By the uniqueness of the classical solution, Prop. 1, it follows that *y* = *y*_∗_ and *z* = *z*_∗_. □

## Notes

Contributing authors;

### Competing Interest Statement

The authors have declared no competing interest.

### Summary of Updates

The new version has the following additions: 1. A slightly different title and abstract. 2. We have edited the text, in particular we have added analytical results for well-posedness and convergence of the method. 3. We have added a new set of simulations.

## References

[1] Lucy V Hiscox, Matthew DJ McGarry, Hillary Schwarb, Elijah EW Van Houten, Ryan T Pohlig, Neil Roberts, Graham R Huesmann, Agnieszka Z Burzynska, Bradley P Sutton, Charles H Hillman, et al. Standard-space atlas of the viscoelastic properties of the human brain. Human brain mapping, 41(18):5282–5300, 2020.

[2] Tomokazu Sato, Mikhail G Shapiro, and Doris Y Tsao. Ultrasonic neuromodulation causes widespread cortical activation via an indirect auditory mechanism. Neuron, 98(5):1031–1041, 2018.

[3] Joseph Blackmore, Shamit Shrivastava, Jerome Sallet, Chris R Butler, and Robin O Cleveland. Ultrasound neuromodulation: a review of results, mechanisms and safety. Ultrasound in medicine & biology, 45(7):1509–1536, 2019.

[4] Claire Rabut, Sangjin Yoo, Robert C Hurt, Zhiyang Jin, Hongyi Li, Hongsun Guo, Bill Ling, and Mikhail G Shapiro. Ultrasound technologies for imaging and modulating neural activity. Neuron, 108(1):93–110, 2020.

[5] Hossein Salahshoor, Mikhail G Shapiro, and Michael Ortiz. Transcranial focused ultrasound generates skull-conducted shear waves: Computational model and implications for neuromodulation. Applied Physics Letters, 117(3):033702, 2020.

[6] Adam Wittek, Karol Miller, Ron Kikinis, and Simon K Warfield. Patient-specific model of brain deformation: Application to medical image registration. Journal of biomechanics, 40(4):919–929, 2007.

[7] Ashutosh Chaturvedi, Christopher R Butson, Scott F Lempka, Scott E Cooper, and Cameron C McIntyre. Patient-specific models of deep brain stimulation: influence of field model complexity on neural activation predictions. Brain stimulation, 3(2):65–77, 2010.

[8] Maxwell Lewis Neal and Roy Kerckhoffs. Current progress in patient-specific modeling. Briefings in bioinformatics, 11(1):111–126, 2010.

[9] Claudia Errico, Juliette Pierre, Sophie Pezet, Yann Desailly, Zsolt Lenkei, Olivier Couture, and Mickael Tanter. Ultrafast ultrasound localization microscopy for deep super-resolution vascular imaging. Nature, 527(7579):499–502, 2015.

[10] Anders M Dale, Arthur K Liu, Bruce R Fischl, Randy L Buckner, John W Belliveau, Jeffrey D Lewine, and Eric Halgren. Dynamic statistical parametric mapping: combining fmri and meg for high-resolution imaging of cortical activity. neuron, 26(1):55–67, 2000.

[11] Simon Chatelin, André Constantinesco, and Rémy Willinger. Fifty years of brain tissue mechanical testing: from in vitro to in vivo investigations. Biorheology, 47(5-6):255–276, 2010.

[12] Silvia Budday, Timothy C Ovaert, Gerhard A Holzapfel, Paul Steinmann, and Ellen Kuhl. Fifty shades of brain: a review on the mechanical testing and modeling of brain tissue. Archives of Computational Methods in Engineering, 27(4):1187–1230, 2020.

[13] R Muthupillai, DJ Lomas, PJ Rossman, James F Greenleaf, Armando Manduca, and Richard Lorne Ehman. Magnetic resonance elastography by direct visualization of propagating acoustic strain waves. science, 269(5232):1854–1857, 1995.

[14] Philip V Bayly, Ahmed Alshareef, Andrew K Knutsen, Kshitiz Upadhyay, Ruth J Okamoto, Aaron Carass, John A Butman, Dzung L Pham, Jerry L Prince, KT Ramesh, et al. Mr imaging of human brain mechanics in vivo: new measurements to facilitate the development of computational models of brain injury. Annals of biomedical engineering, 49(10):2677–2692, 2021.

[15] K. Xu, A. M. Tartakovsky, J. Burghardt, and E. Darve. Learning viscoelasticity models from indirect data using deep neural networks. Computer Methods in Applied Mechanics and Engineering, 387:114124, 2021.

[16] Hossein Salahshoor and Michael Ortiz. Model-free data-driven viscoelasticity in the frequency domain. Computer Methods in Applied Mechanics and Engineering, 403:115657, 2023.

[17] Roderic Lakes. Viscoelastic Materials. Cambridge University Press, 2009.

[18] Sudeshna Patra, Pulickel M Ajayan, and Tharangattu N Narayanan. Dynamic mechanical analysis in materials science: The novice’s tale. Oxford Open Materials Science, 1(1):itaa001, 2021.

[19] EG Herbert, WC Oliver, and GM Pharr. Nanoindentation and the dynamic characterization of viscoelastic solids. Journal of physics D: applied physics, 41(7):074021, 2008.

[20] Erik G Herbert, WC Oliver, A Lumsdaine, and George Mathews Pharr. Measuring the constitutive behavior of viscoelastic solids in the time and frequency domain using flat punch nanoindentation. Journal of materials research, 24(3):626–637, 2009.

[21] PV Bayly, PG Massouros, E Christoforou, A Sabet, and GM Genin. Magnetic resonance measurement of transient shear wave propagation in a viscoelastic gel cylinder. Journal of the Mechanics and Physics of Solids, 56(5):2036–2049, 2008.

[22] Kristy B Arbogast and Susan S Margulies. Material characterization of the brainstem from oscillatory shear tests. Journal of biomechanics, 31(9):801–807, 1998.

[23] R. Abraham, J.E. Marsden, and T. Ratiu. Manifolds, Tensor Analysis and Applications, volume 75 of Applied Mathematical Sciences. Springer, New York, NY, 1988.

[24] Stefan Müller and Michael Ortiz. On the γ-convergence of discrete dynamics and variational integrators. Journal of Nonlinear Science, 14(3):279–296, 2004.

[25] S. Conti, F. Hoffmann, and M. Ortiz. Convergence rates for ansatz-free data-driven inference in physically constrained problems. ZAMM - Journal of Applied Mathematics and Mechanics / Zeitschrift für Angewandte Mathematik und Mechanik, page e202200481, 2023.

[26] Armando Manduca, Philip V Bayly, Richard L Ehman, Arunark Kolipaka, Thomas J Royston, Ingolf Sack, Ralph Sinkus, and Bernard E Van Beers. Mr elastography: Principles, guidelines, and terminology. Magnetic resonance in medicine, 85(5):2377–2390, 2021.

[27] Joseph A Sebastian, Eric M Strohm, Jérôme Baranger, Olivier Villemain, Michael C Kolios, and Craig A Simmons. Assessing engineered tissues and biomaterials using ultrasound imaging: In vitro and in vivo applications. Biomaterials, page 122054, 2023.

[28] Johannes Weickenmeier, Mehmet Kurt, Efe Ozkaya, Max Wintermark, Kim Butts Pauly, and Ellen Kuhl. Magnetic resonance elastography of the brain: a comparison between pigs and humans. Journal of the mechanical behavior of biomedical materials, 77:702–710, 2018.

[29] Manzil Zaheer, Satwik Kottur, Siamak Ravanbakhsh, Barnabas Poczos, Russ R Salakhutdinov, and Alexander J Smola. Deep sets. In I. Guyon, U. Von Luxburg, S. Bengio, H. Wallach, R. Fergus, S. Vishwanathan, and R. Garnett, editors, Advances in Neural Information Processing Systems, volume 30. Curran Associates, Inc., 2017.

